# Redefining the specificity of phosphoinositide-binding by human PH domain-containing proteins

**DOI:** 10.1101/2020.06.20.163253

**Authors:** Nilmani Singh, Adriana Reyes-Ordoñez, Michael A. Compagnone, Jesus F. Moreno Castillo, Benjamin J. Leslie, Taekjip Ha, Jie Chen

## Abstract

Pleckstrin homology (PH) domains are presumed to bind phosphoinositides (PIPs), but specific interaction with and regulation by PIPs for most PH domain-containing proteins are unclear. Here we employed a single-molecule pulldown assay to study interactions of lipid vesicles with full-length proteins in mammalian whole cell lysates. Of 67 human PH domaincontaining proteins examined, 36 (54%) were found to have affinity for PIPs with various specificity, the majority of which had not been reported before. Further investigation of ARHGEF3 revealed structural requirements and functional relevance of its newly discovered PI(4,5)P_2_ binding. A recursive-learning algorithm based on the assay results of the 67 proteins was generated to analyze the sequences of 246 human PH domains, which predicted 48% of them to bind PIPs. A collection of the predicted PIP-binding proteins was assayed, with the vast majority found to bind PIPs. Collectively, our findings reveal unexpected lipid-binding specificity of PH domain-containing proteins.

## INTRODUCTION

Specific phospholipid-protein interactions are critical to the regulation of many signal transduction processes and cellular functions. These interactions typically involve lipid-binding domains recognized by specific lipid species and/or physical properties of the membrane such as charge or curvature (1–4). The largest family of putative lipid-binding domains (LBDs) is the pleckstrin homology (PH) domain, with over 250 members encoded by the human genome (Pfam database). Originally defined by its presence in the protein Pleckstrin (5,6), this domain of 100-120 amino acids has an invariable structure of seven-stranded β-sandwich lined by a C-terminal a-helix (7). PH domains are found in many different types of proteins that are involved in regulating diverse signaling pathways and functions. Some of the earliest characterized PH domains, including those in Pleckstrin, RasGAP, and GRK2, were found to have affinity for PI(4,5)P_2_ (8). The PH domains in PLCδand AKT1 bind with high selectivity to PI(4,5)P_2_ and PI(3,4,5)P3 (PIP3), respectively (9,10). Indeed, the interaction with PIP is so specific for those two PH domains that they have been commonly used as sensors to detect PI(4,5)P_2_ and PIP3 in cells (11,12).

Despite those early examples of PH-PIP interactions, to date only a modest number of PH domain-containing proteins have been demonstrated to bind PIPs with specificity. A comprehensive analysis of *S. cerevisiae* PH domains revealed that only one of them had specific affinity for a particular PIP and the rest of them displayed little affinity or selectivity for PIPs (13). Another study examined a large number of mouse PH domains and found 20% of them to translocate to the plasma membrane (PM) in response to PIP3 production in cells, but most of those PIP3-responsive PH domains did not show specific binding to PIP3 in lipid binding assays (14). Thus, it has been proposed that the specific PIP recognition by PLCδ-PH and AKT1-PH may be the exception rather than the rule for PH domains and that only a small percentage of all PH domains bind PIPs with high affinity and specificity (4). Protein partners have been identified for PH domains, which may cooperate with lipid-PH interaction or operate independently to regulate or mediate PH domain functions (1,15).

It is important to note that our knowledge to date of the lipid binding properties of PH domain-containing proteins is largely derived from studies of isolated PH domains rather than full-length proteins, at least partly owing to the hurdle of purifying full-length proteins for lipid binding assays. Intra-molecular interaction is a universal mechanism for determining protein structure and activity, and this crucial determinant would be eliminated when a PH domain is taken out of the context of the protein. To circumvent the laborious process of purifying proteins and to preserve post-translational modifications of the proteins, we have developed a total internal reflection fluorescence (TIRF) microscopy-based single-molecule pulldown (SiMPull) assay to study protein-lipid interactions with whole cell lysates and lipid vesicles (16), herein referred to as lipid-SiMPull or SiMPull. The sensitivity and specificity of this assay have been demonstrated through highly selective pulldown of various LBDs, as well as full-length AKT1, from cell lysates by vesicles containing PIPs known to interact with those proteins (16). We have now applied the lipid-SiMPull assay to interrogate 67 human PH domain proteins for their binding to vesicles of various compositions. Our results suggest that PIP recognition by PH domain proteins is more prevalent than previously believed. We have also used the assay data to generate a recursive-learning algorithm, which reveals PH domain sequence determinants for PIP binding and predicts PIP binding for the entire family of human PH domains.

## RESULTS

### Lipid-SiMPull assay for human PH domain proteins

In order to investigate binding of PIPs by PH domain proteins using the lipid-SiMPull assay (16), we created cDNA constructs and transfected them in human embryonic kidney (HEK) 293 cells to express 67 full-length human PH domain proteins with EGFP fused at their C-termini. Those 67 proteins were randomly collected based on availability of intact cDNA clones from the hORFeome V5.1 library. Each cDNA in the final plasmid was sequence-validated in its entirety, and expression of the full-length fusion protein in HEK293 cells was confirmed by western analysis (see examples in Supplementary Figure 1A). Concentrations of EGFP-fusion proteins in the lysates were determined by fluorescence intensity and calculated using a standard emission (ex488/em520) curve generated with purified recombinant EGFP (Supplementary Figure 1B). Small unilamellar vesicles (SUVs) were made with 60 mol % phosphatidylcholine (PC), 15 mol % phosphatidylserine (PS), 20 mol % cholesterol, and 5 mol % of one of seven PIPs: PI(3)P, PI(4)P, PI(5)P, PI(3,4)P_2_, PI(4,5)P_2_, PI(3,5)P_2_, and PI(3,4,5)P3. Vesicles containing 5% phosphatidic acid (PA), a different type of signaling lipid than PIPs, were also included in our assays. As a negative control, vesicles were made to include an additional 5 mol % PC in place of PIP (referred to as PC or control vesicles hereafter). All vesicles also had 0.01 mol % biotinylated phosphatidylethanol (PE) to facilitate immobilization on imaging slides.

SiMPull assays were performed with the SUVs immobilized on slides and lysates containing 5 nM EGFP-fusion protein flowed into the slide chamber followed by TIRF imaging (Figure 1A). Each of the 67 proteins was assayed against each of the 9 types of vesicles. The SUVs also contained a lipophilic red dye, DiD, to allow visualization of the immobilized vesicles on slides and confirmation of consistent vesicle density across assays. Pulldown of EGFP proteins was quantified by counting green fluorescent spots. Each preparation of SUV was validated by binding to a positive control in SiMPull, and only the experiments with validated SUVs were further processed. Each vesicle-protein pair was assayed using validated vesicles with at least three independent lysates. Based on characterization of many positive and negative controls with the SiMPull assay, we arrived at 100 EGFP spots per area of 1600 μm^2^ (after background subtraction) as the threshold for binding and the data were interpreted as a binary outcome - binding or no binding. As shown in Supplementary Table 1, the assay results from three independent experiments were remarkably consistent for the vast majority of the 67 proteins.

**Figure 1.**
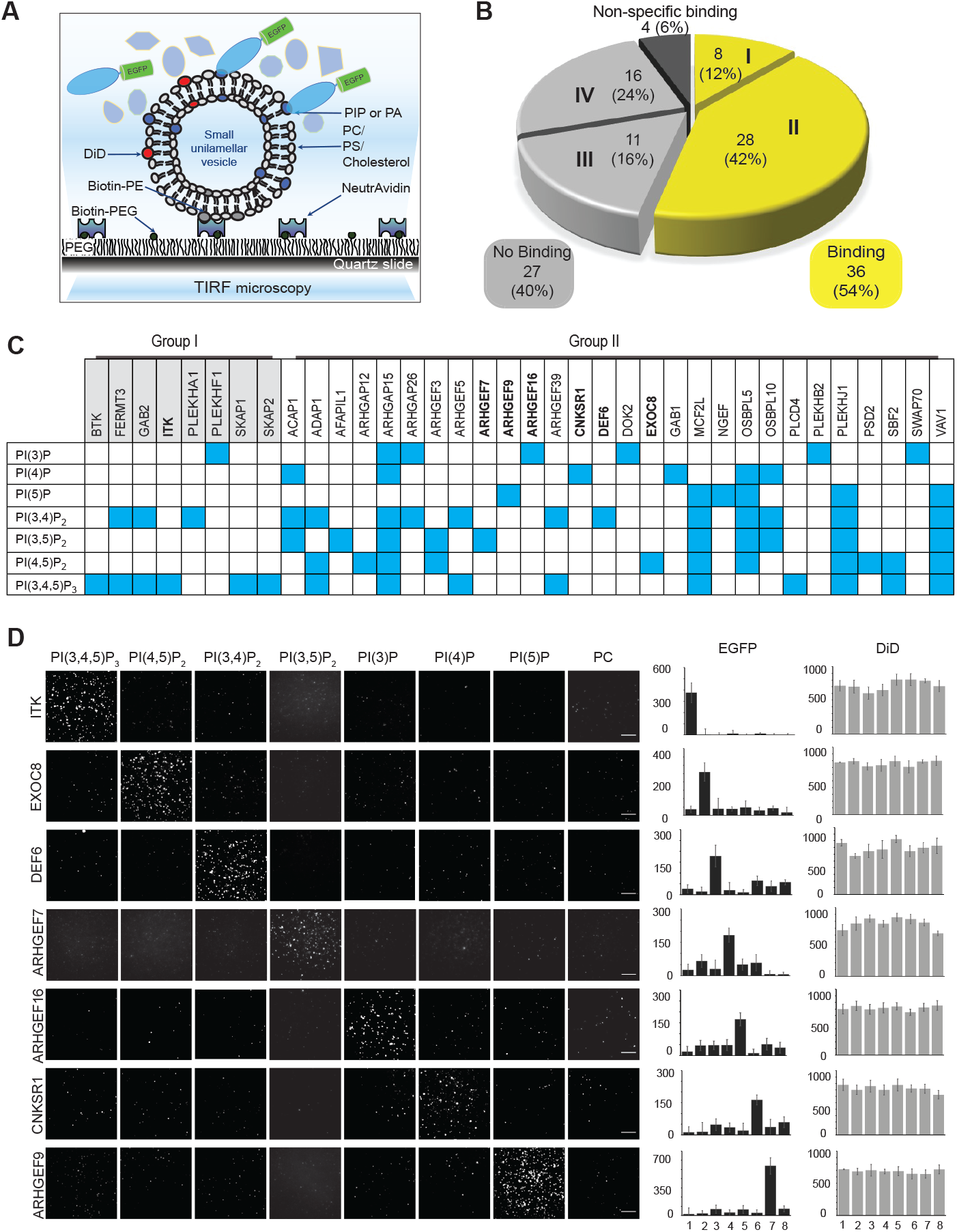
Specific PIP binding by full-length human PH domain proteins. **(A)** Schematic representation of the lipid-SiMPull assay to detect lipid-protein interactions. **(B)** A pie chart summarizing the results of PIP-binding assays for 67 full-length human PH domain proteins against 9 types of vesicles (see text for details). Group I: reported binding confirmed. Group II: novel binding. Group III: binding reported but not found by SiMPull. Group IV: binding not reported nor found by SiMPull. **(C)** A binary representation of results for the 36 proteins found to bind PIPs in SiMPull. The vesicles are identified by the unique PIP, but all contained PC, PS, cholesterol, DiD, and biotin-PE. Blue: binding; white: no binding. SiMPull data for the proteins in bold are shown in D. **(D)** Representative TIRF images of 7 EGFP-fusions pulled down by vesicles containing 5% PIP, with PC as a negative control. EGFP and DiD spots were quantified from 10 or more image areas to yield the average number of spots per 1600 μm^2^ for each assay, shown in the graphs on the right. Each assay was repeated with lysates from at least three independent transfections, and the data can be found in Supplementary Table 1. The results for 67 proteins are summarized in Table 1. Scale bars: 5 μm.

### New PH domain protein-PIP interactions are revealed

The assay results for all proteins are summarized in Table 1, and the raw data can be found in Supplementary Table 1. Of the 67 proteins examined, 4 proteins displayed promiscuous binding to lipids including PA and PC: CERT1, NET1, PLEKHO1 and SPATA13. Misfolding could not be ruled out for those proteins. Thirty-two proteins were found to bind PIPs with various selectivity, and none of them bound PA. Among the 31 proteins that did not bind lipids in our initial assays, some had been previously reported to bind PIPs as either PH domain alone or full-length protein (Supplementary Table 2). We considered the possibility that some of those proteins might require more than 5% PIP in the vesicle for effective binding, since PIPs in the cell membrane can cluster and their asymmetric distribution may lead to drastically higher local concentrations (17). Thus, we performed SiMPull assays with vesicles containing 20% PIP3 or PI(4,5)P_2_ for 10 proteins that had been reported to bind either PIP but not found to bind in our assays (ARHGAP12, DOK1, FERMT2, GRB14, GRK2, OSBPL8, PLCD4, PLEK, SBF2, SKAP_2_). Four of the proteins (ARHGAP12, PLCD4, SBF2, SKAP_2_) indeed bound 20% PIP3 or PI(4,5)P_2_.

In total, 36 of the 67 proteins (54%) were found to bind PIPs with some specificity (Table 1, Figure 1B). The positive binding data are also illustrated in a binary table in Figure 1C. Representative SiMPull assay images and quantification are shown in Figure 1D for 7 proteins, each displaying specificity for a different PIP. Of the 36 proteins found to bind PIPs, our assay results confirmed reported PIP binding by 8 proteins (Group I in Figure 1B & 1C), whereas 28 proteins (Group II) had been either not previously reported to bind any lipid or reported to display different lipid binding profiles than those found in our assays.

Of the 27 proteins found not to bind PIP in our assays, 11 had been reported to bind lipids (Group III) and 16 not reported (Group IV). It is possible that those 11 represent false negatives in our assay. Previously we showed that the assay detected LBD-lipid interactions that were reported to have K_d_’s in the high nanomolar/sub-micromolar range (16). To estimate the threshold of detection under our assay conditions, we performed SiMPull with the PX domain of p40Phox, the WT, R60A, and K92A mutant of which bind PI(3)P with K_d_ of 5 μM, 17.5 μM, and >50 μM, respectively (18). As shown in Supplementary Figure 2, robust binding to PI(3)P, but not to PI(4,5)P_2_, was detected for WT p40Phox-PX. Binding to the R60A mutant was also observed, at just above the cut-off of 100 EGFP spots, whereas the K92A mutant did not bind the lipid. Therefore, our assay under current conditions has a detection threshold of affinity in the 10-20 μM range. It is unlikely that many physiologically relevant interactions would have been missed.

### Phosphatidylserine is not responsible for the novel PH domain protein-PIP interactions

Phospholipids may cooperate in the membrane to determine specific interaction with proteins (2,3). The presence of sphingolipids and PS in the membrane has been found to contribute to enhanced affinity and/or specificity for PIPs by yeast PH domains (19). Since the SUVs in our assays contained 15 mol% PS (based on mammalian cell membrane composition), we wondered whether the newly discovered PIP-PH protein interactions were dependent on PS. To address this possibility, we re-assayed 23 of the new PIP-binding proteins (Group II) with SUVs containing only PC and a specific PIP, omitting PS and cholesterol. Remarkably, all 23 proteins bound their respective PIPs without PS in the vesicles (Figure 2A). Representative SiMPull images are presented in Figure 2B. Therefore, the presence of PS does not appear to account for the novel PIP-PH protein interactions emerged from our studies.

**Figure 2.**
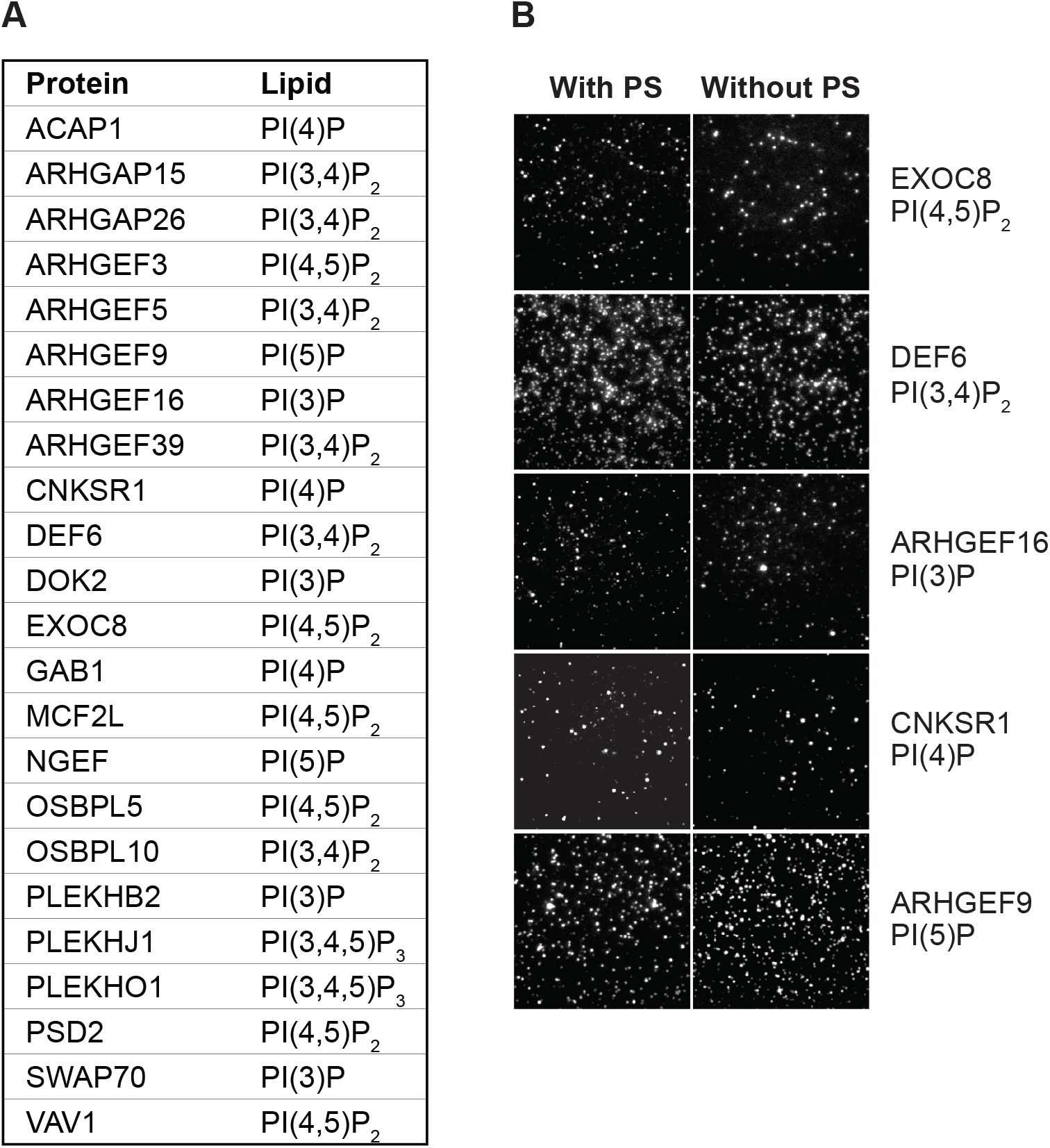
The presence of PS in lipid vesicles does not affect PIP binding by proteins. **(A)** SiMPull was performed for 23 proteins against lipid vesicles containing the indicated PIPs, with or without PS and cholesterol. Binding was observed for all with both types of vesicles. For each condition at least two independent experiments were performed with similar results. Data were analyzed as described for Table 1 and Supplementary Table 1. **(B)** Representative TIRF images of EGFP pulldown for 5 proteins that are also shown in Figure 1D, each by vesicles with or without PS.

### PH domain of ARHGEF3 is necessary for its specific lipid binding

To compare the behaviors of PH domains to their full-length protein counterparts, we performed SiMPull assays with EGFP fusions of 12 PH domains from 10 proteins found to bind PIP in our assays (two of them have two PH domains each). To our surprise, with the exception of PLEKHA1, the PH domains either did not bind PIP or bound different PIPs than their fulllength counterparts (Supplementary Table 3). This discrepancy could be attributed to misfolding of the PH domains when expressed alone and/or interference by the EGFP tag.

We further examined the PIP binding by ARHGEF3 (also named XPLN for exchange factor found in platelets, leukemic, and neuronal tissues), a GEF for RhoA and RhoB (20). ARHGEF3 (Figure 3A) belongs to the Dbl family of 70 human RhoGEFs, the vast majority of which contain a Dbl homology (DH) domain and a PH domain in tandem (21). There had been no report of specific lipid binding by ARHGEF3. In our assay, the full-length ARHGEF3 protein bound specifically to PI(4,5)P_2_ and PI(3,5)P_2_ (Figure 3B). However, the PH domain alone fused with EGFP at either the N- or C-terminus did not bind PI(4,5)P_2_ vesicles, neither did fragments of ARHGEF3 missing the N-terminal 125 amino acids (ΔN) or the PH domain (ΔPH) (Figure 3C). Expression of the various ARHGEF3 proteins was confirmed by western blotting shown in Supplementary Figure 3A. Hence, it appears that the entire full-length ARHGEF3 may be necessary to confer lipid binding.

**Figure 3.**
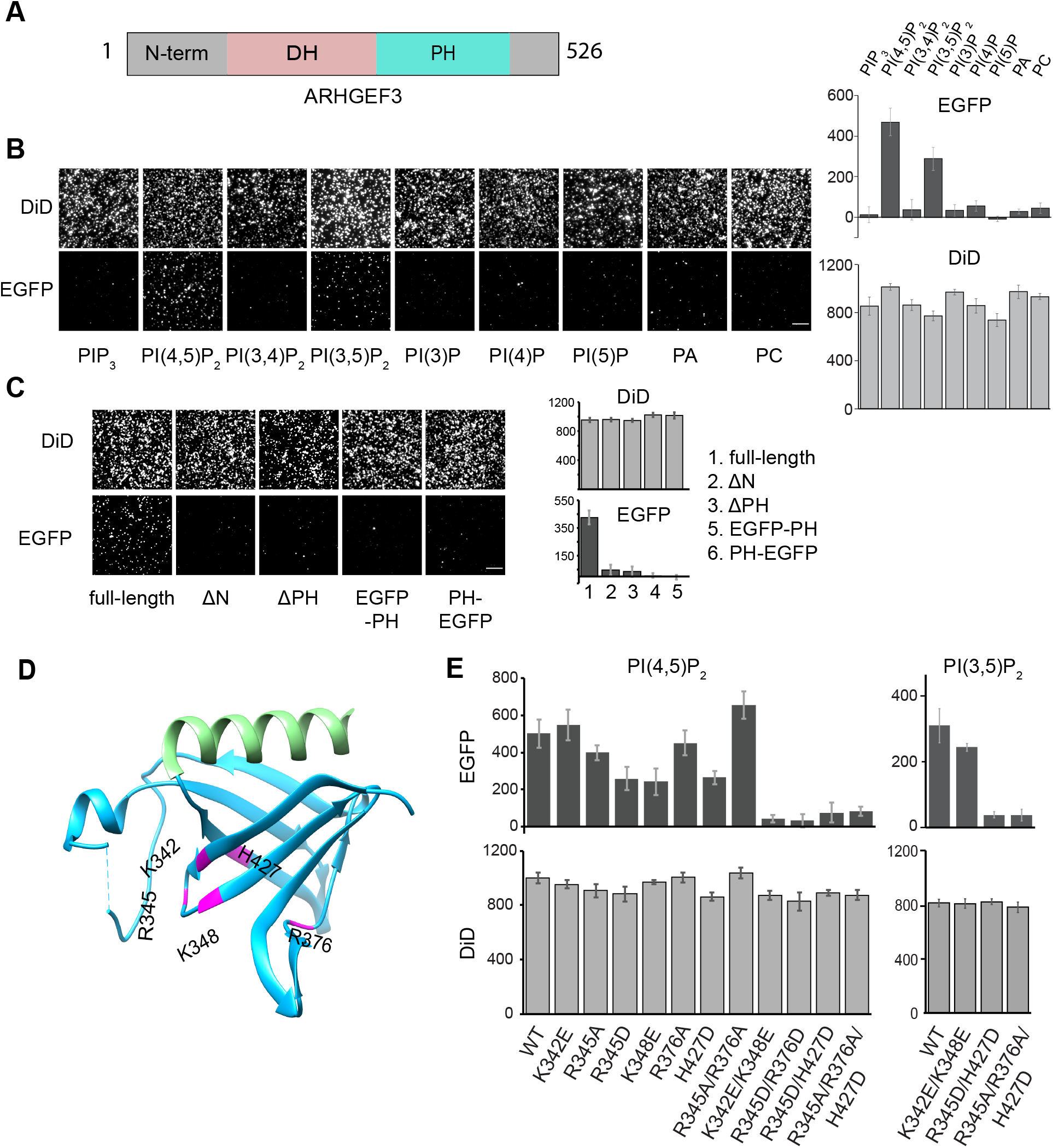
ARHGEF3 binds PI(4,5)P_2_ through its PH domain. **(A)** Schematic diagram of ARHGEF3 domain structure. **(B)** HEK293 cells were transiently transfected with EGFP-ARHGEF3, and the lysates were subjected to SiMPull assays against various types of lipid vesicles as indicated. Representative TIRF images are shown. EGFP and DiD spots were quantified from 10 or more image areas to yield the average number of spots per 1600 μm^2^ for each assay, shown in the graphs on the right. Each assay was performed with at least 3 independent transfections with similar results. **(C)** Similar to B, various EGFP-tagged ARHGEF3 fragments were assayed for binding to vesicles containing 5% PI(4,5)P_2_. **(D)** Ribbon representation of the structure of ARHGEF3 PH domain (PDB 2Z0Q), with residues predicted to interact with PIP highlighted in magenta. **(E)** Similar to B, EGFP-tagged ARHGEF3 mutants were assayed for binding to 5% PI(4,5)P_2_ or PI(3,5)P_2_ vesicles. Each assay was repeated with lysates from at least 3 independent transfections. Scale bars: 5 μm.

As another approach to assess whether the PH domain of ARHGEF3 was required for interaction with PIP, we considered structural information available in the literature. In reported PH domain-PIP headgroup interactions, negatively charged amino acids in the β1-β2 and β3-β4 loops play critical roles (1). Applying the Patch Finder Plus software (22) to the reported ARHGEF3-PH crystal structure (23), we identified a positively charged patch containing K342, R345, K348, R376 and H427 to be potentially involved in the interaction between ARHGEF3 and PIP (Figure 3D). These amino acids were mutated to alanine or acidic residues in full-length ARHGEF3 individually and in combination, and the mutant proteins (Supplementary Figure 3A&B) were subjected to SiMPull assays with PI(4,5)P_2_ vesicles. Mutation of any one of the positively charged residues did not result in drastic reduction of PI(4,5)P_2_ binding, but three double-mutants (K342E/K348E, R345D/R376D, R345D/H427D) and a triple-mutant (R345A/R376A/H427D) each abolished lipid binding by ARHGEF3 (Figure 3E). Therefore, despite the requirement of the entire protein for PI(4,5)P_2_ binding, the PH domain of ARHGEF3 most likely contributes directly to the PIP-protein interaction.

Next, we investigated PI(3,5)P_2_ binding by the ARHGEF3 mutants. As shown in Figure 3E, the triple mutant and R345D/H427D abolished PI(3,5)P_2_ binding. Interestingly, the K342E/K348E mutant retained PI(3,5)P_2_ binding activity. These observations suggest that the PH domain of ARHGEF3 may interact with the two PIPs via structurally distinct mechanisms. Future investigation of these molecular interactions would be informative.

### PI(4,5)P_2_ binding is necessary for ARHGEF3 PM targeting, activation of RhoA, and induction of stress fiber formation

Because PI(4,5)P_2_ is highly abundant in the PM, we wondered if ARHGEF3 binding to PI(4,5)P_2_ played a role in the protein’s membrane recruitment. Recombinant EGFP-ARHGEF3 was found throughout the cell including the nucleus, and the nuclear localization is consistent with the presence of a nuclear localization sequence in ARHGEF3. We observed prominent presence of EGFP-ARHGEF3 in the PM of NIH3T3 cells and, importantly, the PI(4,5)P_2_ binding-deficient mutant of ARHGEF3, K342E/K348E, displayed drastically reduced localization to the PM (Figure 4A). This observation is consistent with ARHGEF3 targeting to the PM via its interaction with PI(4,5)P_2_.

**Figure 4.**
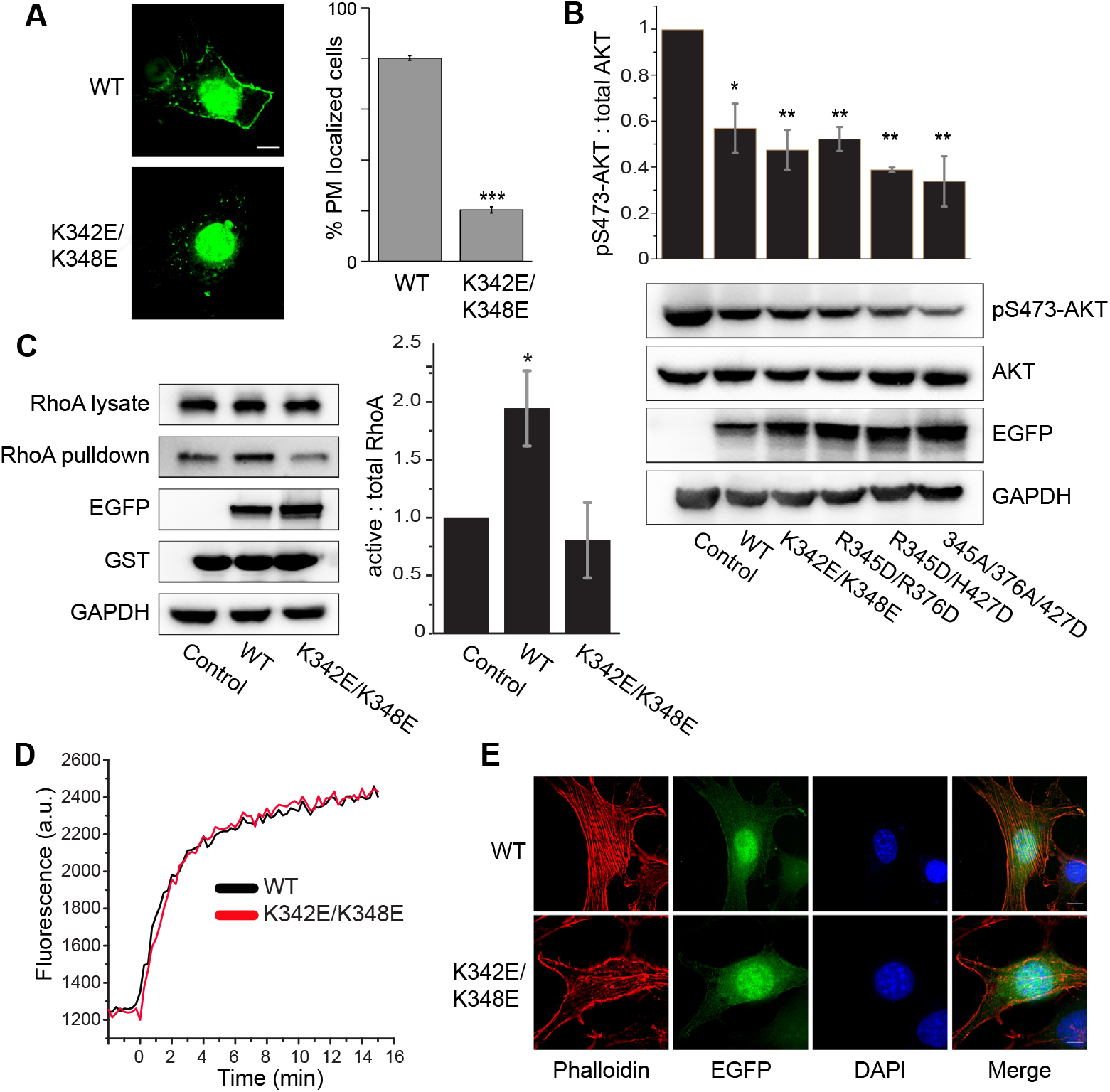
PI(4,5)P_2_ binding is necessary for ARHGEF3 membrane targeting, cellular activity and function. **(A)** NIH3T3 cells were transfected for 24 hrs with EGFP-tagged wild-type or lipid binding-deficient mutant K342E/K348E of ARHGEF3. The transfected cells were then treated with 0.03% digitonin for 2.5 min to remove cytosolic proteins before fixation. Percentage of transfected cells with PM localization was quantified from 3 independent experiments. **(B)** HEK293 cells were transfected with wild-type or lipid binding-deficient mutants of ARHGEF3, serum-starved overnight, and then stimulated with 10% FBS for 30 min followed by cell lysis and western blotting. Western blot signals were quantified by densitometry to generate ratio of pAKT versus total AKT from 3 independent experiments. **(C)** HEK293 cells were transfected for 24 hrs with empty vector (control), wildtype, or K342E/K348E mutant of ARHGEF3. Cell lysates were subjected to active RhoA pulldown assay using GST-rhotekin beads, and analyzed by western blotting. Western blot signals from 5 independent experiments were quantified by densitometry to generate the relative ratios of RhoA pulled down (active) and RhoA in lysates (total) shown in the graph. **(D)** Purified wild-type and K342E/K348E mutant ARHGEF3 were subjected to in vitro RhoA guanine nucleotide exchange assay. Three independent experiments were performed with similar outcome, and representative results are shown. **(E)** NIH3T3 cells were transfected for 24 hrs with wild-type or K342E/K348E mutant of ARHGEF3, followed by fixation and phalloidin/DAPI staining. Three independent experiments were performed with similar outcome, and representative images are shown. Error bars represent SD. Student’s t test was performed to compare WT and mutant in A, and each sample to the control in B & C. *\*P*< 0.05; *\*\*P*< 0.01; *\*\*\*P*< 0.001. Scale bars: 5 μm.

Next, we asked whether PI(4,5)P_2_ binding is relevant to ARHGEF3’s function in a cell. ARHGEF3 has two unrelated activities: as a GEF for RhoA/B (20) and as a GEF-independent inhibitor of mTORC2 phosphorylation of AKT (24). We examined these activities of ARHGEF3 in HEK293 cells. As shown in Figure 4B, expression of wild-type ARHGEF3 suppressed AKT phosphorylation as expected, and expression of each lipid binding-deficient mutant of ARHGEF3 (K342E/K348E, R345D/R376D, R345D/H427D, or R345A/R376A/H427D) was equally effective in inhibiting AKT, suggesting that PI(4,5)P_2_ binding is not involved in ARHGEF3’s function as an inhibitor of mTORC2 (nor is PI(3,5)P_2_ binding). This is consistent with the previous observation that the N-terminal 125-amino acid fragment, devoid of the PH domain, is sufficient to inhibit mTORC2 (24). On the other hand, unlike the wild-type protein, the K342E/K348E mutant ARHGEF3 was ineffective in activating cellular RhoA activity upon overexpression, as measured by pulldown of active RhoA using an effector protein (Figure 4C). Hence, lipid binding is apparently necessary for ARHGEF3’s GEF activity toward RhoA in the cell.

We also performed in vitro GEF assays with ARHGEF3 and ARHGEF3-K342E/K348E proteins purified from bacteria, and found that the two proteins displayed near identical activities towards RhoA (Figure 4D). We were not able to assess the effect of lipids on the GEF activity because we found addition of lipid vesicles to be incompatible with the GEF assay. Nevertheless, our observation confirmed that the K342E/K348E mutant retained proper folding and enzymatic activity. The simplest explanation for all our results combined is that PI(4,5)P_2_ binding and/or PM targeting is necessary to achieve maximum GEF activity for ARHGEF3 in the cell.

Consistent with its role as a GEF for RhoA, ARHGEF3 has been reported to regulate actin cytoskeleton reorganization and assemble stress fibers in the cell (20). To probe a potential role of ARHGEF3-PI(4,5)P_2_ interaction in this process, we expressed EGFP-ARHGEF3 in NIH3T3 cells and visualized actin cytoskeleton with phalloidin staining. As shown in Figure 4E, expression of wild-type ARHGEF3, but not the K342E/K348E mutant, led to robust stress fiber formation, suggesting that PI(4,5)P_2_ binding may be critical for ARHGEF3 regulation of actin cytoskeleton reassembly, a RhoA-dependent process. Taken together, our observations uncover a novel mechanism of ARHGEF3 regulation and provide functional validation of a protein-lipid interaction identified in our lipid-SiMPull assays.

### A recursive-learning algorithm for probabilistic sequence comparison based on SiMPull data

Although our survey of 67 proteins covered only 1/4 of the PH domain-containing protein family in the human genome, we reasoned that our results could have predictive power for the rest of the family. Given the high likelihood that the PH domains mediate PIP binding of the full-length proteins, we compared the sequences of the PH domains of the 67 proteins we studied plus AKT1-PH as a postive control, excluding the 4 proteins that bound lipids non-specifically. We also excluded two proteins (ADAP1 and AFAP1L1) that each contained two PH domains and bound PIP as full-length proteins, because we did not know which PH domain contributed to the binding. This yielded a collection of 67 PH domains. Based on reported structural studies of the PH domain-PIP headgroup interaction, the loop between the first two βstrands (β1-β2) with a basic sequence motif of “KX_n_(K/R)XR” plays the most prominent role (3). However, we found the vast majority of the 67 PH domains to contain this motif, with no distinction between those that bound PIPs and those that did not (Figure 5A). Therefore, we next considered the possibility that sequence features throughout the entire PH domain are determinants of PIP binding, as suggested by Park et al. for PIP3-regulated PH domains (14).

**Figure 5.**
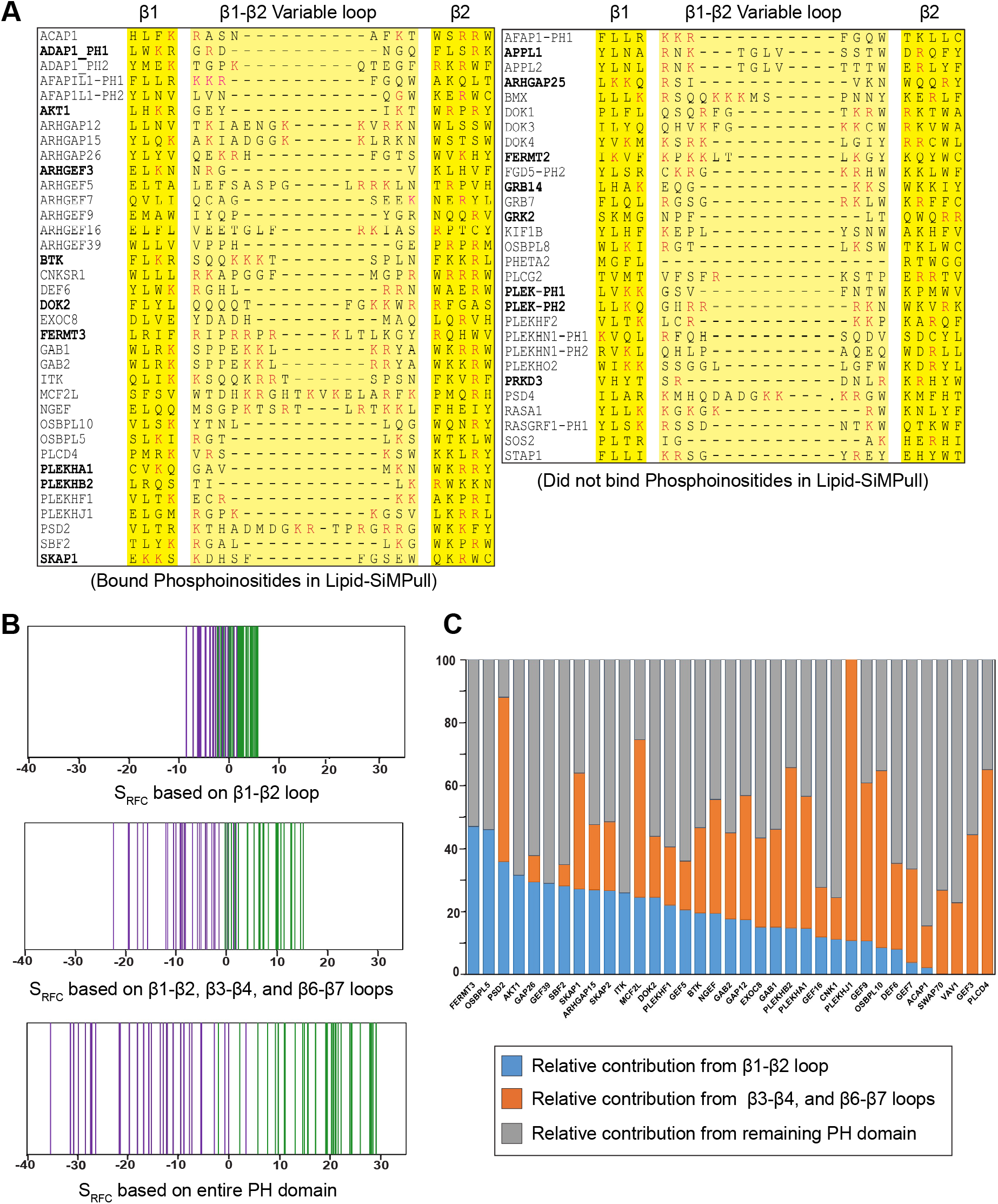
Recursive-learning algorithm distinguishes PIP-binding when taking the entire PH domain into consideration. (**A**) Sequence alignment of PIP-binding (left) and non-binding (right) PH domains in the β1- β2 region. Bold: PH domains with reported crystal structures. Red: residues conforming to the KXn(K/R)XR motif. Three PH domains (AFAP1-PH2, FGD5-PH1 and RASGRF1-PH2) were not shown in the alignment due to an unusually short β1 sheet. (**B**) S_RFC_ scores were calculated for the 67 PH domains in our assays, with amino acids from β1- β2 variable loop, β 1-β2, β3-β4, β6-β7 variable loops, or the entire PH domain. Each vertical line represents one PH domain. Green: bound PIP in SiMPull assays; purple: did not bind PIP in SiMPull assays. (**C**) Relative contribution by β 1-β2 loop, β3-β4 and β6-β7 loops, or the rest of the PH domain is plotted based on S_RFC_ for each protein that bound PIP in SiMPull assays.

To identify sequence signatures of PIP binding, we employed an unbiased probabilistic sequence comparison strategy developed by Park et al., called recursive functional classification (RFC) (14). This strategy relies on accurate sequence alignement of multiple PH domains, which can be a challenge given the low sequence similarity shared by PH domains despite the 3D structure conservation. We employed the PROMALS3D (<u>PRO</u>file <u>M</u>ultiple <u>A</u>lignment with predicted <u>L</u>ocal <u>S</u>tructures and <u>3D</u> constraints) tool, which makes use of available 3D structural information to guide multiple sequence alignments (25), to align the sequences of 246 PH domains from 222 human proteins in the Pfam and UniProt databases, including the 67 PH domains described above. We found a total of ~280 PH domains in the database, but excluded ~30 of them from the alignment due to their unusual sequence features that would have caused very large gaps in the alignment. Alignment of the 246 PH domains yielded 297 amino acid positions; the 67 PH domains from proteins investigated in SiMPull and the other 179 PH domains are listed in Supplementary Table 4 and Supplementary Table 5, respectively. Following the reported method (14) we created two probability matrices of 297 (positions) x 20 (amino acids): P^B^ for the PIP binding group (35 PH domains including AKT1-PH), and P^NB^ for the non-binding group (32 PH domains). Based on the assumption that each sequence position contributes independently to PIP binding, we created an RFC-matrix, with each element of the matrix calculated as log(P^B^/P^NB^). In the RFC-matrix values > 0 represent positive correlation with PIP binding whereas values < 0 represent negative correlation. This RFC-matrix can be used to calculate the predicted PIP binding score (S_RFC_) for any PH domain, as previously described (14).

### Recursive functional classification identifies potential amino acid determinants of PIP binding distributed throughout the entire PH domain

Using the RFC-matrix created above, we first scored positions in the β1-β2 loop and its flanking regions to calculate S_RFC_. The resulting scores separated PIP-binding proteins from nonbinding proteins without a large margin (Figure 5B). When we expanded the positions to include β3-β4 and β6-β7 loops and their flanking regions, which are believed to also contribute to PH-PIP interactions (1), the separation of the two groups of proteins improved (Figure 5B). However, when all positions of the PH domain were taken into calculation, the PIP-binding and non-binding groups were even more cleanly separated with markedly higher scores in both the positive and negative ranges (Figure 5B). This outcome validates the idea that the entire PH domain may be involved in determining whether a domain (or full-length protein) binds PIP. Interestingly, the relative contributions of various regions of the PH domain to PIP-binding vary drastically among those found to bind PIPs (Figure 5C), suggesting that the structural basis of PH-PIP interactions may be diverse within this family of proteins.

We created a heat map (Figure 6A) to represent the RFC-matrix for the 67 PH domains studied in SiMPull and aligned it to the sequence of AKT1-PH based on its crystal structure (PDB ID: 1UNP). Amino acids with high impact on lipid binding, either positively or negatively, were distributed throughout the PH domain. Following the reported method (14), we determined the overall contribution of each amino acid position to PIP binding, with 12 positions of the highest contribution to PIP binding marked (Figure 6B). While some of those residues are located in the classically defined head group binding pocket, many are not. Other than those 12, additional positions spanning the entire PH domain are likely to also play important roles in PIP binding (Figure 6B). It should be noted that this is a composite view, and that an individual PH domain would use only a subset of those residues for interaction with PIP (see Figure 5C).

**Figure 6.**
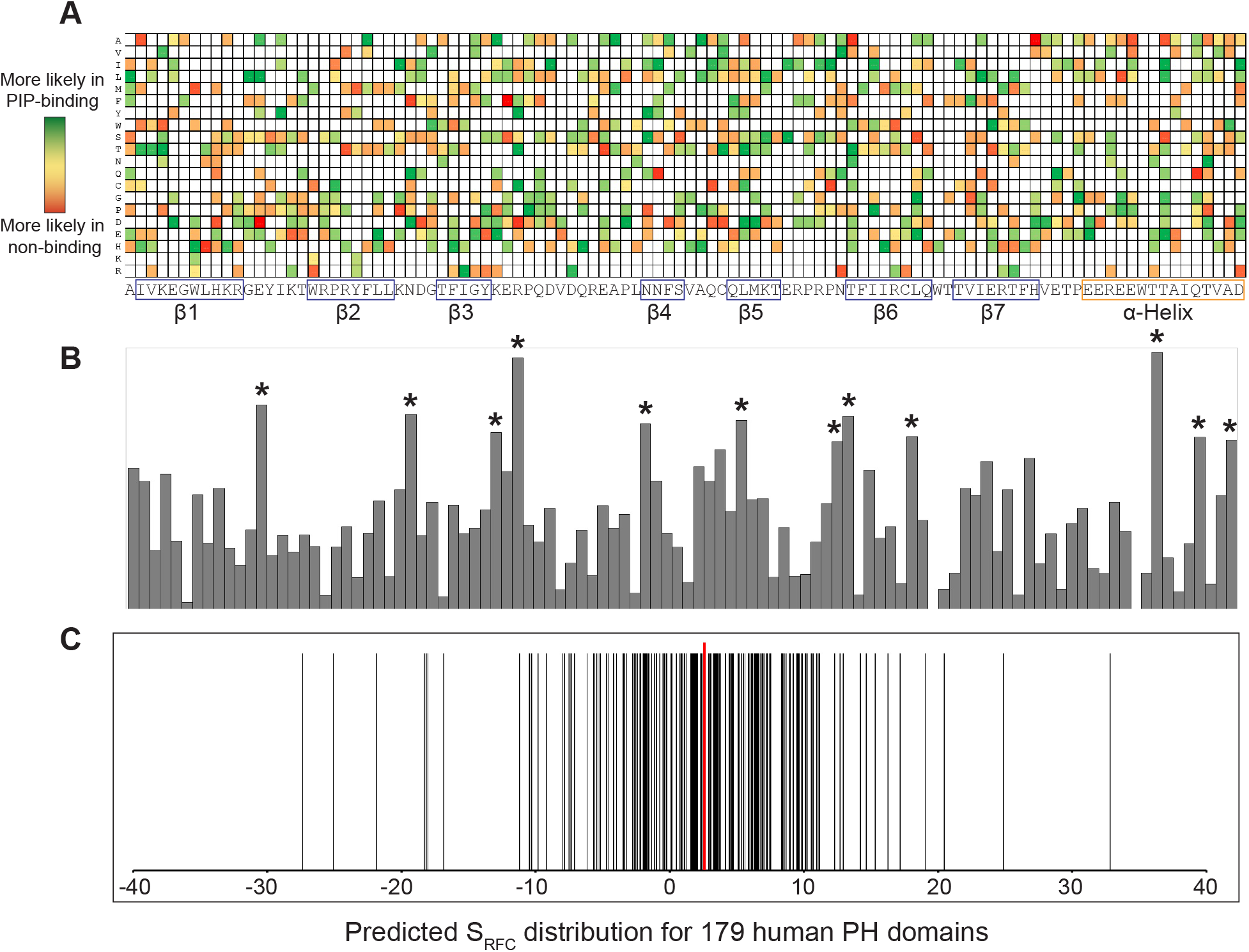
Recursive-learning algorithm identifies amino acid determinants of PIP-binding and predicts binding by human PH domains. (**A**) A heat map showing the contribution of 20 amino acids at each position for the 67 PH domains. The amino acid positions are aligned to AKT1-PH domain based on its crystal structure (PDB ID: 1UNP). (**B**) Contribution from different positions of PH domain towards PIP-binding was plotted from RFC-matrix. Twelve positions with the highest scores are marked by stars and aligned to the sequence in A. (**C**) Prediction of PIP-binding for 179 human PH domains using the RFC-matrix. Each vertical black line represents one PH domain. The red line indicates the cutoff value (2.59) for PIPbinding.

### The RFC algorithm predicts 50% of human PH domain-containing proteins to bind PIP

Next, we applied the RFC algorithm obtained above to analyze the human PH domains not investigated by SiMPull. While in principle any protein with a positive S_RFC_ score can be considered to possibly bind PIP, we set out to establish a more stringent cut-off score for positive correlation with PIP binding by scrambling the sequences of all 246 PH domains and subjecting them to scoring by the same algorithm (Supplementary Figure 4). Using mean + 3x SD, we arrived at 2.59 as the threshold S_RFC_ score for a PH domain to be considered likely to bind PIP. Of the 179 PH domains not tested by SiMPull, 86 (48%) scored above the threshold (Figure 6C and Supplementary Table 5). Taking into consideration our assay data, the overall estimate is that 50% of the PH domains in the human genome bind PIPs.

To test the validity of the RFC algorithm, we performed SiMPull assays with predicted binders that have not been reported to bind any lipids to date. The availability of cDNA clones and confirmation of EGFP-fusion full-length protein expression led to the random selection of 20 proteins in that category (Supplementary Table 6). We first confirmed that none of the 20 proteins displayed nonspecific binding to PC. Because our goal was to validate PIP binding and the binding to any PIP would satisfy the criterion, we assayed the 20 proteins against one of the three PIPs frequently found to bind PH domains: PIP3, PI(3,4)P_2_, and PI(4,5)P_2_. If binding was confirmed for one PIP, that protein was not assayed against other PIPs. As shown in Supplementary Table 6, 19 out of the 20 proteins were found to bind a PIP. The only exception was PSD3, which did not bind any of the three PIPs although it is still possible that it binds to a PIP not yet tested. These results validate the predictive power of our recursively learning approach and, furthermore, are consistent with the notion that the PH domain defines the observed PIP binding.

## DISCUSSION

Our analysis of a random collection of 67 human PH domain-containing proteins using the lipid-SiMPull assay reveals that 54% of them bind PIPs and that many of the specific interactions have not been reported before. As an example of functional validation of the novel lipid-protein interaction emerged from our assays, we have demonstrated that PI(4,5)P_2_ binding is necessary for the GEF activity of ARHGEF3 in the cell. A recursive-learning algorithm based on the results of the 67 proteins has predicted 48% of the remaining human PH domains in the family to bind PIPs with some specificity. The predictive power of this algorithm is confirmed by our finding that 19 out of 20 proteins predicted to bind PIP (but never reported to bind lipids) indeed display affinity for at least one PIP in SiMPull assays. Our work suggests that ~50% of all human PH domain proteins likely bind PIPs with specificity, in contrast to the previous estimate that only 10% of all PH domains bind PIPs with high affinity and specificity (4).

The SiMPull assay offers a unique advantage of using full-length proteins expressed in the native environment (i.e., mammalian cells) without the need for purification. These proteins would be likely to maintain their native structures and proper post-translational modifications. However, due to overexpression of the EGFP-fusion proteins it is unlikely that any endogenous proteins in the lysate would reach stoichiometric levels to contribute directly to the observed recombinant protein binding to lipid vesicles. Hence, the lipid-protein interactions observed in lipid-SiMPull assays are most likely intrinsic properties of the PH domain-containing proteins. Our experimental validation of the PIP binding predicted by a recursive learning algorithm based on PH domain sequences also suggests that the PH domain defines PIP binding by the full-length proteins.

We have determined the threshold of detection under the current lipid-SiMPull assay conditions to be in the K_d_ range of 10-20 μM, which is sufficiently sensitive to detect most reported and/or physiologically relevant protein-PIP interactions. Although the possibility of interference by the EGFP tag has not been ruled out for those proteins that did not display any lipid binding in our assays, we do not consider it likely that many true interactions have been missed by our assay. Instead, we would like to suggest that non-physiological binding conditions (such as in the lipid strips assay) and/or isolation of the PH domains out of the context of fulllength proteins may have been accountable for at least some of the discrepancies between our observations and those in the literature. Indeed, of those 11 proteins that were reported to bind PIPs but found not to bind in our assays (Group III), 10 were based on isolated PH domains and/or lipid strips assays (references in Supplementary Table 2).

The PIP binding specificity discovered in our study can potentially lead to new understanding of the functions and mechanisms of the PH domain proteins. For instance, previously the PH domain of CNKSR1 (or CNK1) had been found to bind PIPs weakly and non-specifically (26), but in our assays the full-length CNKSR1 protein bound specifically to PI(4)P. Interestingly, CNKSR1 has been reported to promote PI(4,5)P_2_ production from PI(4)P by phosphatidylinositol 4-phosphate 5-kinases (PIP5Ks) at the PM where insulin receptor signaling takes place (27), although the mechanism is unclear. Based on our finding, it is conceivable that CNKSR1 is targeted to PI(4)P-enriched membrane in order to activate PIP5K or to recruit PIP5K to its substrate. Another example is EXOC8 (or Exo84), a component of the exocyst, which is an octameric protein complex involved in tethering secretory vesicles to the plasma membrane for fusion during exocytosis (28). A generally accepted view is that Sec3 and Exo70 in the exocyst bind PI(4,5)P_2_ and target the entire octameric complex to the membrane. Our observation of PI(4,5)P_2_ binding by EXOC8 suggests that EXOC8 may also contribute directly to the regulation of the exocyst by PI(4,5)P_2_, and that further investigation could potentially lead to revision of the current model of exocyst assembly.

The vast majority of the 70 members of the Dbl family of RhoGEFs contain a PH domain immediately following the catalytic DH domain (21). Although those PH domains had been speculated to bind phospholipids and subsequently contribute to membrane localization and/or allosteric regulation of the RhoGEFs, very few of them have been reported to have marked affinity for specific PIPs. Thus, the idea of phospholipids regulating RhoGEFs through their PH domains remains controversial (29). Among the proteins we studied with the SiMPull assay, 14 are RhoGEFs belonging to the Dbl family. PIP binding was observed for 9 of them (ARHGEF3, ARHGEF5, ARHGEF7, ARHGEF9, ARHGEF16, ARHGEF39, MCF2L, NGEF, VAV1), all of which were either never reported to bind lipids or reported to bind lipids with different specificity than what we observed. For instance, the PH domain of ARHGEF3 was reported to bind phospholipids with little selectivity in lipid strips assays (14). In contrast, we have found that full-length ARHGEF3 binds PI(4,5)P_2_ and PI(3,5)P_2_, and that the binding is dependent on the PH domain. Interestingly, a mutant (K342E/K348E) ARHGEF3 has lost PI(4,5)P_2_ binding while retaining its binding of PI(3,5)P_2_, thus offering an ideal tool to establish the importance of PI(4,5)P_2_ binding for the GEF activity of ARHGEF3. Future investigation is warranted to probe a role of PI(3,5)P_2_ in ARHGEF3 function. Follow-up studies on the 8 remaining RhoGEFs and investigation of lipid binding by other Dbl family members will also likely be illuminating.

The recursive-learning algorithm developed by Park et al. (14) is an unbiased probabilistic sequence comparison strategy that is now facilitated by accurate multiple sequence alignment incorporating 3D structural information (25). The clean separation of PIP binding and nonbinding groups of PH domains by this algorithm and, more importantly, our validation of predicted novel PIP binding, attest to the power of the method. Our RFC analysis suggests that for some proteins PIP binding may involve regions of the PH domain outside of the previously known interacting site. Similar suggestions have been made by others for PIP3 regulation (14) and organelle PIP binding (19) by PH domains. Interestingly, two most recent molecular dynamics simulation studies of the PH domain of GRP1 have revealed that there may be multiple PIP3 binding sites – canonical and non-canonical – on this domain (30,31). Future biochemical and structural studies to probe these putative interactions of novel modes will be informative.

## MATERIALS AND METHODS

### Cell Culture

HEK293 cells were maintained in high-glucose DMEM with 10% FBS, 2 mM L-glutamine, and penicillin/streptomycin at 37 ^o^C in 5% CO2. NIH3T3 cells were maintained in high-glucose DMEM with 10% newborn calf serum and 4 mM L-glutamine, and penicillin/streptomycin at 37 ^o^C in 5% CO2. For SiMPull experiments, HEK293 cells were transfected for 24 hours in 6-well plates using PolyFect® (3 μL/μg DNA) following manufacturer’s recommendations. For immunofluorescence experiments, cells were plated in 12- well plate on poly-L-lysine coated coverslips and transfected with Lipofectamine^TM^ 3000 following manufacturer’s recommendations. For active RhoA pulldown assays, HEK293 cells were transfected in 10-cm plates with Lipofectamine™ 3000.

### Cell Lysis and Western Blotting

For SiMPull assays, the cells were collected after 24-hr transfection in detergent-free buffer (40 mM HEPES, pH8.0, 150 mM NaCl, 10 mM β-glycerophosphate, 10 mM sodium pyrophosphate, 2 mM EDTA, 1x Sigma protease inhibitor cocktail). The cells were lysed by sonication for 3 seconds on ice followed by ultracentrifugation at 90,000xg in a TLA100.3 rotor for 1 hour at 4°C. For SiMPull, EGFP concentration was measured using a standard emission (ex488/em520) curve of pure recombinant EGFP, and each cell lysate was diluted in vesicle buffer (10 mM Tris·HCl, pH 8.0, 150 mM NaCl) to yield 5 nM EGFP. For western blotting, cells were lysed in 1x SDS sample buffer or as described above and mixed at 1:1 with 2x SDS sample buffer, both containing β-mercaptoethanol at a final concentration of 5%, and heated for 5 min at 95 ^o^C. Proteins were resolved by SDS-PAGE and transferred onto PVDF membrane. The membrane was incubated with primary and secondary antibodies following manufacturers’ recommendations. HRP-conjugated secondary antibody was reacted with West Pico PLUS Chemiluminescent Substrate, and the signal was detected using an Invitrogen iBright digital imager.

### cDNA Cloning and Mutagenesis

Plasmids harboring cDNAs for human PH domain-containing proteins from human cDNA library hORFeome V5.1 were cloned as a pool into the pDest-eEGFP-N1 vector using Gateway Cloning, followed by identification of individual clones. Additional cDNA clones were obtained from DNASU, and cloned individually into the pcDNA3-EGFP vector using Gibson assembly. The PH domains were cloned into the pEGFP-C1 vector using Gibson assembly. ARHGEF3 (XPLN) expression plasmids for full-length and truncations were previously reported (24). p40PhoxPX-EGFP was obtained from Addgene (#19010) (32). All point mutants were created using site-directed mutagenesis using QuikChange Lightning Site-Directed Mutagenesis Kit following the manufacturer’s protocol. All cDNAs in the final plasmids were sequence-confirmed in their entirety.

### Lipid Vesicle Preparation

All lipids were mixed in chloroform (0.166 μmol total) and dried under nitrogen flow. The dried mixture was re-suspended in 100 μL vesicle buffer (10 mM Tris·HCl, pH 8.0, 150 mM NaCl) to a final concentration of 1.66 mM. After 30 min incubation at room temperature, vesicles were formed by water bath sonication (Laboratory Supplies, Hicksville, NY, model G112SPIT, 600 v, 80 kc, and 0.5 A) in 4 cycles of 4 min each. Small unilamellar vesicles were collected as the supernatant after ultracentrifugation at 194,398xg in a TLA100.3 rotor for 1 hr at 25 ^o^C.

### Single-Molecule Pulldown Assay

Quartz slides were prepared as described in Jain et al. and Arauz et al. (16,33). Briefly, the slides were thoroughly cleaned and passivated with PEG doped with 0.1-0.2% biotin-PEG. Neutravidin (200 μg/mL) was incubated in chambers for 10 min, followed by addition of biotinylated lipid-vesicles. Freshly prepared whole-cell lysate (80 μL) was added by flowing into each slide chamber, replacing the vesicle solution. An inverted total internal reflection fluorescence (TIRF) microscope with Olympus 100x, NA1.4 lens and EMCCD camera (Andor iXon Ultra 897) was used to acquire single-molecule data at 10 frames/second. Diode pumped solid state lasers were used to excite EGFP at 488nm (Coherent) and DiD at 638nm (Cobalt). All SiMPull experiments were performed at room temperature. Average number of spots per imaging area (~1600 μm^2^) was calculated from 10 or more images.

### SiMPull Image Analysis

TIRF image acquisitions were processed in IDL to generate image files, and the number of fluorescence spots were identified with point-spread function. MATLAB® was used to extract data from data files generated in IDL and import into Excel. The background number of EGFP spots was gathered prior to addition of lysate and subtracted from EGFP spots after lysate addition to generate data for each image. At least 10 SiMPull images (1600 μm^2^ each) were analyzed to generate the average number of spots per imaging area for each assay. The IDL scripts used to process raw image files are publicly available from Taekjip Ha laboratory (http://ha.med.jhmi.edu/resources/).

### Fluorescence Imaging

HEK293 or NIH3T3 cells on poly-L-lysine-coated glass coverslips were fixed in 3.7% paraformaldehyde at room temperature for 15 min and permeabilized with 0.1% Triton X-100 for 5 min. For visualizing stress fibers, Rhodamine phalloidin was diluted 1:3000 and incubated with 3% BSA/PBS at 4 ^o^C for 60 min, followed by incubation with DAPI (1:2500) for 20 min at room temperature. For digitonin permeabilization experiments, cells on coverslips were treated with 0.03% digitonin (20 mM HEPES, PH 7.5, 110 mM KOAc, 5 mM NaOAc, 2 mM Mg(OAc)2, 1 mM EGTA) for 2.5 min, followed by fixation with 3.7% paraformaldehyde. A personal deconvolution microscope system (DeltaVision, Applied Precision) was used with a 100x or 60x NA 1.4 lens to capture fluorescence images. Deconvolution used an enhanced ratio iterative-constrained algorithm (34). The acquired images were processed in ImageJ. For quantification of PM-localized cells, 60-100 transfected cells were counted in each experiment.

### Protein Purification

GST-ARHGEF3 wild-type and K342E/K348E proteins were expressed in E. coli from the pGEX-4T-1 vector and purified using glutathione Sepharose beads following the manufacturer’s recommendations.

### GTPase RhoA Activity Assays

Cellular RhoA activity was measured as previously described (24). Briefly, HEK293 cells in 10-cm plate were transfected for 24 hrs and then lysed in 50 mM Tris (pH 7.4), 10mM MgCl2, 500 mM NaCl, 1% (vol/vol) Triton X-100, 0.1%SDS, 0.5% sodium deoxycholate, and 1x Protease Inhibitors cocktail. The cell lysates were incubated for 1 hr at 4 ^o^C with 60 μg of GST-rhotekin beads, followed by washing with 50 mM Tris (pH 7.4), 10 mM MgCl2, 150 mM NaCl, 1% (vol/vol) Triton X-100, and 1x Protease Inhibitors cocktail, and analysis by western blotting with an anti-RhoA antibody. To measure nucleotide exchange activity of RhoA in vitro, the mant-GTP exchange assay was performed following manufacturer’s recommendation in 20 μL reaction volume in a 384-well plate. In short, purified RhoA (1 μM) was added to exchange buffer (40 mM Tris-HCl, pH 7.5, 100 mM NaCl, 20 mM MgCl2, 4 μM mant-GTP) and fluorescence (ex360/em440) was recorded for 5 min before adding purified GST-ARHGEF3 or GST-ARHGEF3-K342E/K348E.

### RFC Matrix Generation and PIP-binding Prediction

The recursive-learning strategy described by Park et al. (14) was followed, with the details specific to our study described here. The list of human PH domain-containing proteins was obtained from the Pfam database and individual PH domain sequences were acquired from UniProt. After initial alignment of all sequences using PROMALS3D with default parameters, about 30 sequences were removed due to large insertions, and the remaining 246 PH domains were re-aligned. MATLAB was used to create two 297 x 20 probability matrices for PIP-binding (P^B^) and non-binding proteins (P^NB^), respectively. An amino acid was scored in the probability matrix if it was found in at least one PH domain in the group. The RFC matrix was then calculated by taking the log of the ratio of PB to P^NB^. In Figure 5B, the value at each amino acid position is the average of sum-square of values in the corresponding position. For scoring a PH domain, the values in the RFC matrix for matching amino acids at all positions were summed.

## ACKNOWLEDGEMENTS

We thank Dr. Lisa Stubbs and Mr. Joe Troy for help with the initial identification of PH domain sequences, and Ms. Lucy Yao and Ms. Eleanor Marcet for assistance with western blotting. This work was supported by grants from the National Institute of General Medical Sciences (R01 GM089771 to JC and R35 GM122569 to TH).

## AUTHOR CONTRIBUTIONS

Conceptualization and experimental design by N.S., A.R.-O., B.J.L., T.H. and J.C.; experimentation, data collection and analysis by N.S., A.R.-O., M.A.C. and J.F.M.; resources by B.J.L. and T.H.; writing by N.S., A.R.-O., and J.C.; editing by N.S., A.R.-O., T.H. and J.C.; funding acquisition by T.H. and J.C.

---

**Supplementary Figure 1.**
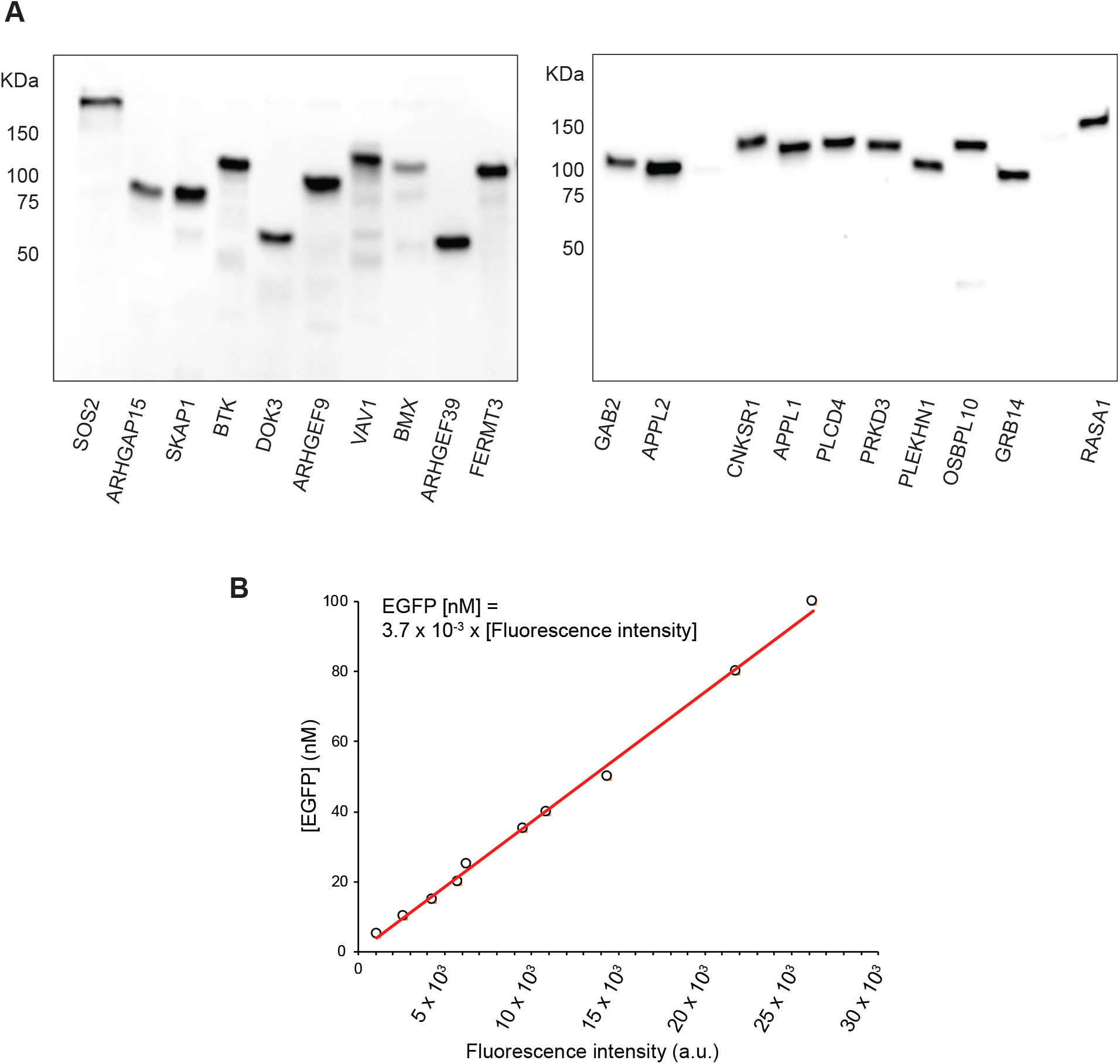
**(A)** Examples of EGFP-fusion proteins transiently expressed in HEK293 cells. Cell lysates were subjected to western blotting with an anti-GFP antibody. **(B)** Fluorescence of pure recombinant EGFP (ex 488 ∕ em 520) was measured at 10 different concentrations on a Spectramax GeminiXPS plate reader (Molecular Devices) to yield a linear standard curve.

**Supplementary Figure 2.**
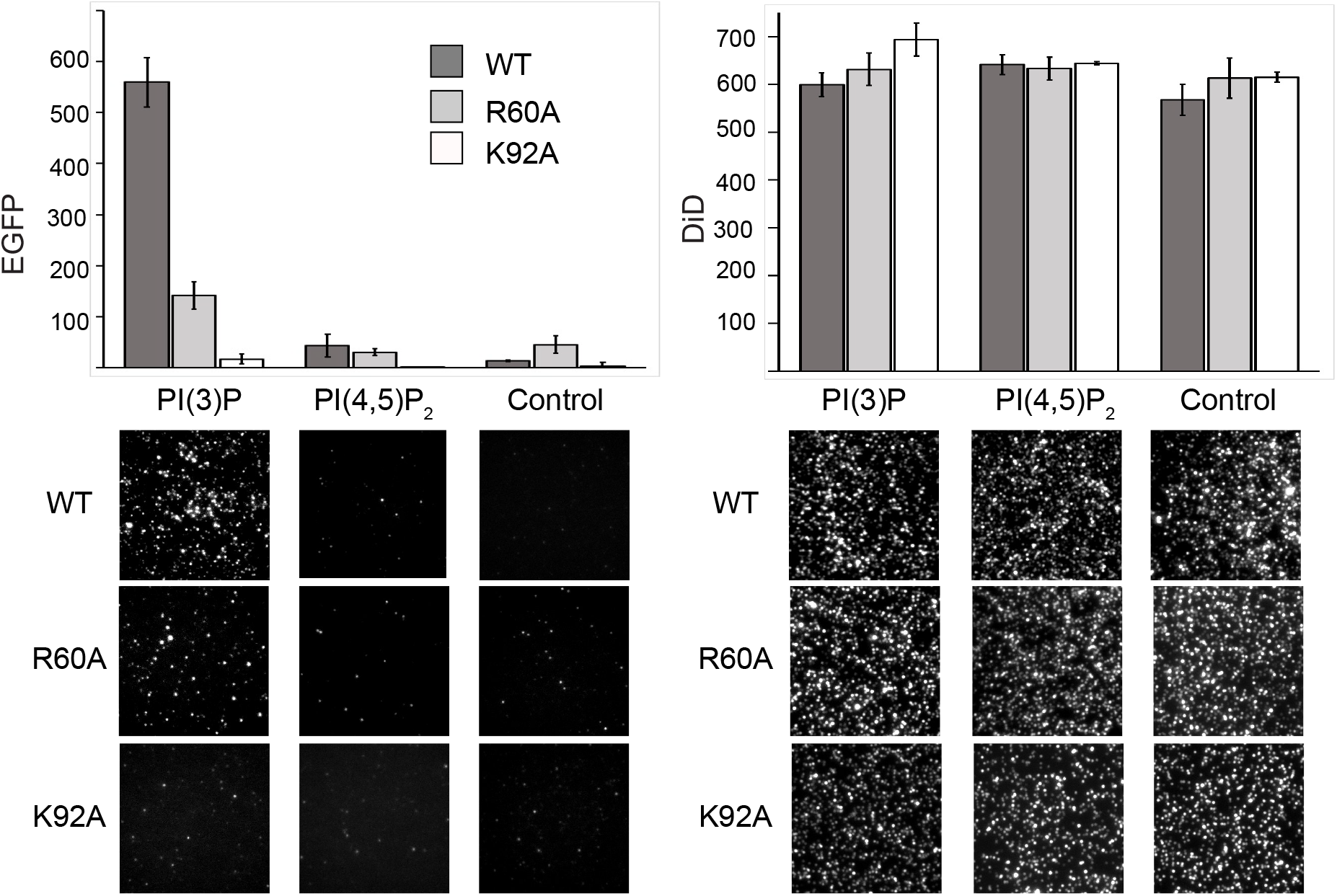
EGFP-p40PhoxPX and mutants were expressed in HEK293 cells, and cell lysates (5 nM EGFP-fusion) were subjected to SiMPull assay with PI(3)P, PI(4,5)P_2_ and PC (control) vesicles. EGFP and DiD (vesicle) spots were quantified as described in Figure 1 legend. The average results of 3 independent experiments are shown with error bars representing SEM.

**Supplementary Figure 3.**
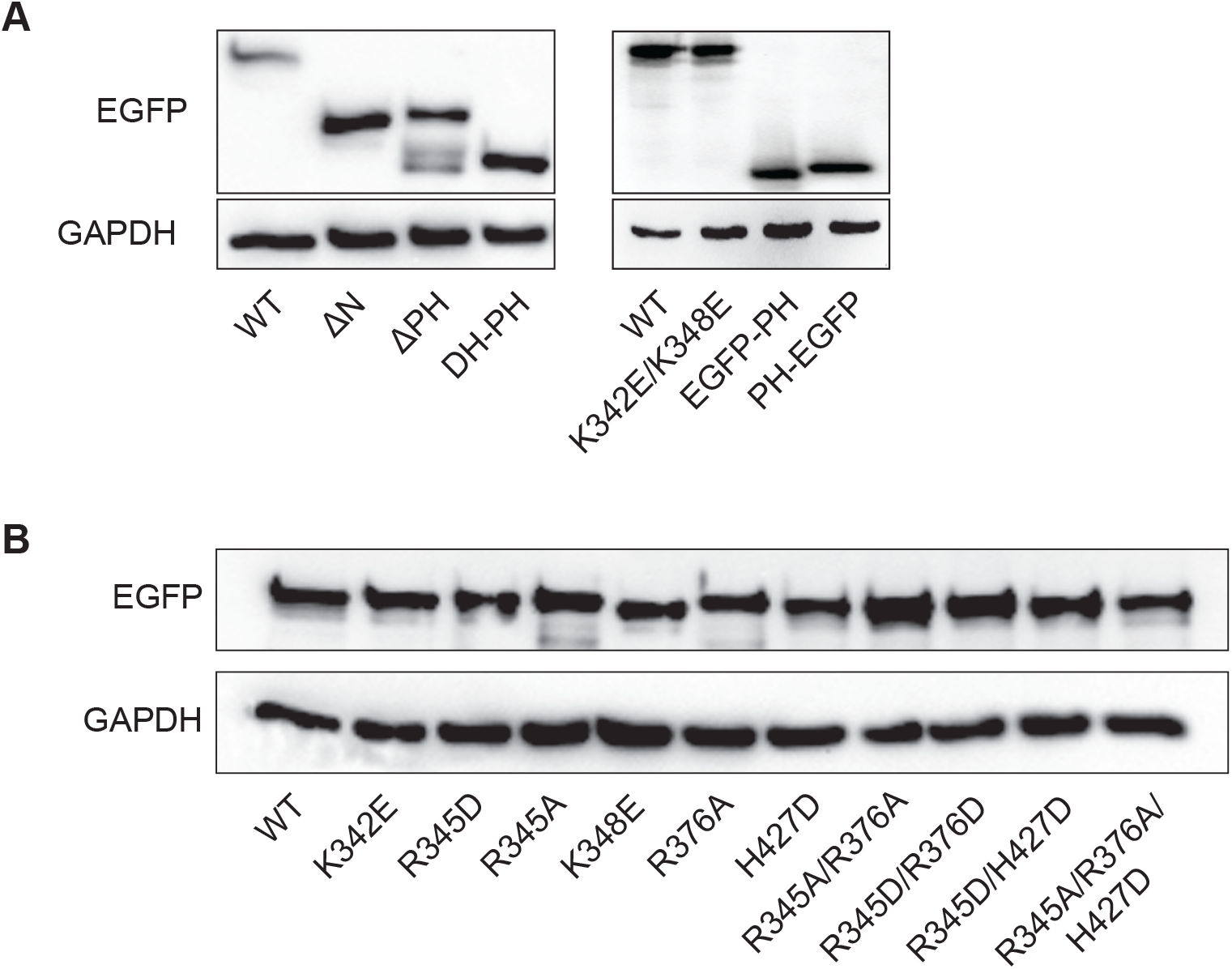
HEK293 cells were transiently transfected to express various truncations **(A)** or point mutants **(B)** of ARHGEF3 fused to EGFP. Cell lysates were analyzed by western blotting with an anti-GFP antibody, and anti-GAPDH blotting served as loading control.

**Supplementary Figure 4.**
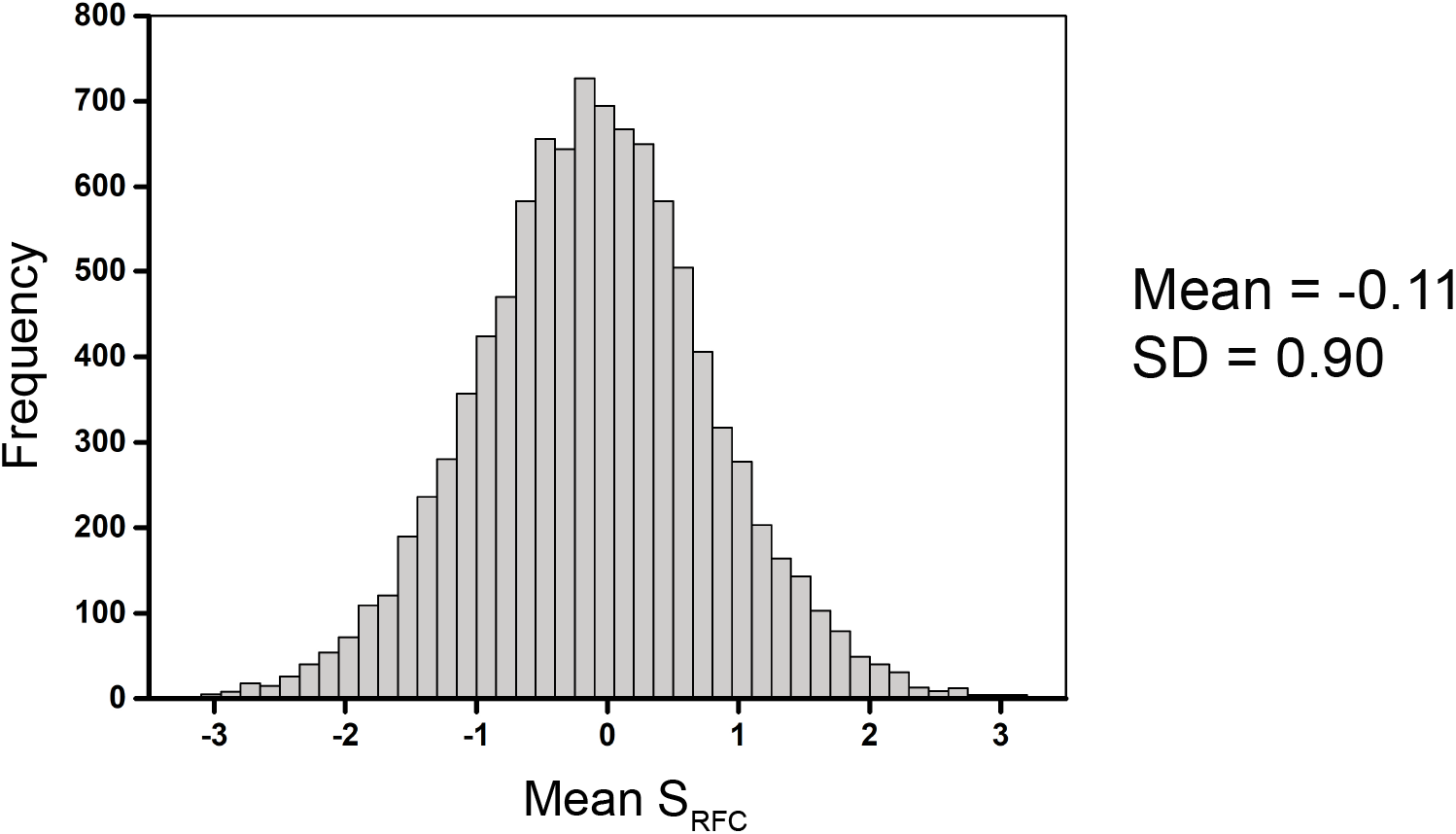
The sequences of 246 human PH domains were scrambled 10,000 times, and the S_RFC_ for each scrambled sequence was calculated. The distribution of mean S_RFC_ is shown, with the overall mean and standard deviation indicated.

