## Supplementary Table 3 for "Redefining the specificity of phosphoinositide-binding by human PH domain-containing proteins"

| **PH Domain** | **Lipid binding by PH domain** | **Lipid binding by full-length protein** |
|---|---|---|
| ACAP1_PH | $PI(3,4)P_2$ | $PI(3,4)P_2$, PI(4)P |
| ADAP1_PH1 | None | $PI(3,4,5)P_3$, $PI(4,5)P_2$, $PI(3,4)P_2$ |
| ADAP1_PH2 | None | $PI(3,4,5)P_3$, $PI(4,5)P_2$, $PI(3,4)P_2$ |
| ARHGEF3-PH | None | $PI(4,5)P_2$, $PI(3,5)P_2$ |
| ARHGEF5_PH | None | $PI(4,5)P_2$, $PI(3,4)P_2$ |
| ARHGEF9_PH | None | PI(5)P |
| DOK2-PH | PI(3,4,5)P3, PI(3)P, PI(4)P | PI(3)P |
| FERMT3-PH | None | $PI(3,4,5)P_3$, $PI(3,4)P_2$ |
| PLEKHA1-PH1 | None | $PI(3,4)P_2$ |
| PLEKHA1-PH2 | $PI(3,4)P_2$ | $PI(3,4)P_2$ |
| SPATA13-PH | None | Bound all lipid vesicles |
| VAV1-PH | None | $PI(3,4,5)P_3$, $PI(4,5)P_2$, $PI(3,4)P_2$, $PI(3,5)P_2$, PI(5)P |

**Supplementary Table 3. SiMPull assay results for selected PH domains.** Experimental details and data processing were as described for Table 1 and Supplementary Table 1. At least three independent experiments were performed with similar outcome for each PH domain.
